# A systematic screen for morphological abnormalities during fission yeast sexual reproduction identifies a mechanism of actin aster formation for cell fusion

**DOI:** 10.1101/103176

**Authors:** Omaya Dudin, Laura Merlini, Felipe Bendezú, Raphaël Groux, Vincent Vincenzetti, Sophie G Martin

## Abstract

In non-motile fungi, sexual reproduction relies on strong morphogenetic changes in response to pheromone signaling. We report here on asystematic screen for morphological abnormalities of the mating process in fission yeast *Schizosaccharomyces pombe*. We derived a homothallic (self-fertile) collection of viable deletions which, upon visual screening, revealed a plethora of phenotypes affecting all stages of the mating process, including cell polarizati cell fusion and sporulation. Cell fusion relies onthe formation of the fusion focus, an aster-like F-actin structure that is marked by strong local accumulation of the myosin V Myo52, which concentrates secretion at the fusion site. A secondary screen for fusion-defective mutants identified the myosin V Myo51-associated coiled-coil proteins Rng8 and Rng9 as critical forthe coalescence of the fusion focus Indeed, *rng8∆* and *rng9∆* mutant cells exhibitmultiple stable dots a the cell-cell contact site, instead of the single cusfo observed in wildtype. Rng8 and Rng9 accumulate on the fusion focus, dependent on Myo51 and tropomyosin Cdc8A. tropomyosin mutant allele, which compromises Rng8/9 localization but not actin binding, similarly leads to multiple stable dots instead of a single focus.By contrast, *myo51* deletion does not strongly affect fusion focus coalescenceWe. propose that focusing of the actinfilaments in the fusionaster primarily relies on Rng8/9-dependent cross-linking of tropomyosin-actin filaments.

## Introduction

Sexual reproduction is carried out by most eukaryotes and permits the alternation of haploid and diploid life stages. It relies on the formation of differentiated haploid cell types that are able to meet and fuse form a zygote, which eventually returns to the haploid state through meiosis. Many of heset events rely on morphological changes, especially in organisms without cell motility. Yeast model systems have been used over decades to uncover basic principles of cell organization, yet no systematic screening of their sexual reproduction process has been performed. Here, we have used the fission yeast *Schizosaccharomyces pombe* to systematically screen for viable gene deletions causing a morphological abnormality in the sexua reproduction process. We anticipated this screen would shed light on the processes of cell polarization, cell-cell fusion and sporulation.

All natural *S. pombe* isolates live as haploid cells, and many, such as the *h90* lab strain, are self-fertile (homothallic) [1, 2]. These cells, which can be of two distinct mating types, P and M, regularly switch mating type by recombination of the silent mating cassette into the active site after cell division, thus resulting ina near genetically identical population that can reproduce sexually [3]. Sexual differentiation is initiated by nitrogen starvation, which leads to the expression of pheromones and cognate receptor on the two cell typ Pheromone signaling involves a GPCR-MAPK signal transduction cascade, which in turn reinforces sexual differentiation and initiates the morphological program of mating [4]. Upon sensing low pheromone levels, cells initially polarize secretion towards a cortical patch assembled around the active form of the small GTPase Cdc42[5]. This patch dynamically forms at various cortical locations and disassembles over time, but cells do not grow. Pheromone secretion and sensing are thought to occur at the patch, which stabilizedis through unknown molecular mechanismsupon higher local pheromone perception, such that two neighboring cells become locked together when their patches meet[6]. Paired cells then grow towards each other to form a pre-zygote, with cell wall still separating the tw partner cells.

To achieve cellfusion, the cell wall needs to be digested at the zone of cell-cell contact to allow plasma membrane fusionThis. relies on the fusion focus, a dedicated actin aster nucleated by the formin-family protein Fus1, which promotes the convergence on a small cortical zone of secretory vesicles transported by type V myosin motors[7–9]. In particular, these motors transport glucanases, enzymes that hydrolyze the bonds linking the cell wall glucan polymer[7]. Over the course of the fusion process, het fusion focus formsfrom an initially broad distribution at the cell projection tip, and stabilizes into a singly focus in opposing locations in the two partner cells. Thistabilization stems from a positive feedback between concentration of pheromone signal at the secretion zone and local enrichment of the pheromone signal transduction machinery, which immobilizes the fusion focus through unknown mechanism [10]. In turn, spatial stabilization permits the focused delivery of glucanases for local cell wall digestion.

Formation of the fusion focus is likely to require several actin-binding proteins, in addition to Fus1. In particular, profilin Cdc3 and tropomyosin Cdc8 are enriched on the structure and necessary f cell-cell fusion [11,12]. Type V myosins also localize on the fusion focus and contribute to its focalization [7, 13]. There are two such myosins in fission yeast[14,15]: Myo52 is the main cargo transporter for both cell polarization and cell fusion[7,16–18], and moves processively on tropomyosin-decorated actin filaments[19]; Myo51 is more unusual, as many of its functions are independent of its cargo-binding tail [7,20,21]. In addition, Myo51 is a single-headed motor protein, and both in vivo and in vitro experiments have shown that only motor ensembles were capable of processive movement[21,22]. In vivo, a dimer of two coiled-coil proteins, Rng8 and Rng9, associates with Myo51, regulates its localization during mitotic growth, and was proposed to contribute to Myo51 processivity by forming higher-order oligomers in vivo[21]. In vitro, the Rng8/9-Myo51 complex was also shown to bind tropomyosin-decorated F-actin independently of the motor domain, thus forming a bivalent-actinF-binding complex cross-linking and sliding actin-tropomyosin filaments relative to one another [22]. Despite these recent advances, how these motors or other actin-binding proteins function to focus an actin aster is not established.

Upon local cell wall digestion, plasma membranes fuse. Though multi-pass transmembrane proteins such as Prm1 have been suggested to participate in this process in several fungal species, the mechanism remains completely elusive[23–24,25]. As the fusion pore then expands the neck connecting the now fused cells is remodeled to create an elongated zygote in which the two parental haploid nuclei fuse. The diploid nucleus then enters meiosis to return the genome to its haploid state, forming four meiotic products, each of which is packaged in a stress-resistant spore. Sporulation is a very morphologically demanding process in which new plasma membrane and new wall is laid down, initiated from the spindle pole associated with each of the four genomic meiotic products[26].

Previous forward-genetic screens have identified a number of steril fusion-defective and sporulation-deficient mutants, and a targeted genome-wide screen for sporulation-defective deletion strains was published in the course of this work [27]. However, there has not been any systematic reverse-genetic screen of the mating process. Here, we present the results of a visual screen for morphological abnormalities during the mating process in fission yeastOur. screen led us to identify the Rng8/9 dimer and its interaction with tropomyosin critical for the formation of the actin fusionfocus. We propose that cross-linking of tropomyosin-actin filaments serves to focalize filaments in the fusion focus.

## Results

### Creation and visual screening of a homothallic deletion collection

To systematically screen the collection of viable deletions for mating defects, we converted the available heterothallic *h+* deletion library [28] to a homothallic *h90* collection by applying a modified version of the SpSGA method [29]. We first integrated a nourseothricin resistance cassette (*natMX*) 6kb away from the expressed *mat1* mating-type cassette, between the genes *mag2* and *rpt6*, in an otherwise wildtype homothallic *h90* strain. Because the genomic region located between the expressed *mat1* locus and silent *mat* loci represents a genetic distance of only 1cM [30], the *h90* trait and *natMX* largely co-segregate, allowing for selection for the *h90* trait at the population level. We then robotically crossed this *h90-natMX* strain to all *kanMX*-marked deletion strains of the *h+* collection in 384-well plate format. Mating was induced on solid medium with low nitrogen for 4 days at 25°CVegetative. haploid cells that had not mated and diploid cells that had not sporulated were killed incubationby at 42°C for 4 days[29]. We note that diploid killing was efficient, as azygotic tetrads, which stem from the sporulation of diploid cells rather than zygotes formed by cell-cell fusion, were observed in only 76/227 strains upon the visual screening described below. Spore germination was triggered by replica plating on solid rich medium (YE). A second replication step to solid medium containing both G418 an nourseothricin selected for homothallic *h90* deletion-carrying strains. Finally, strains were saved at −80°C in YE 25% glycerol (Figure 1A).

**Figure 1:**
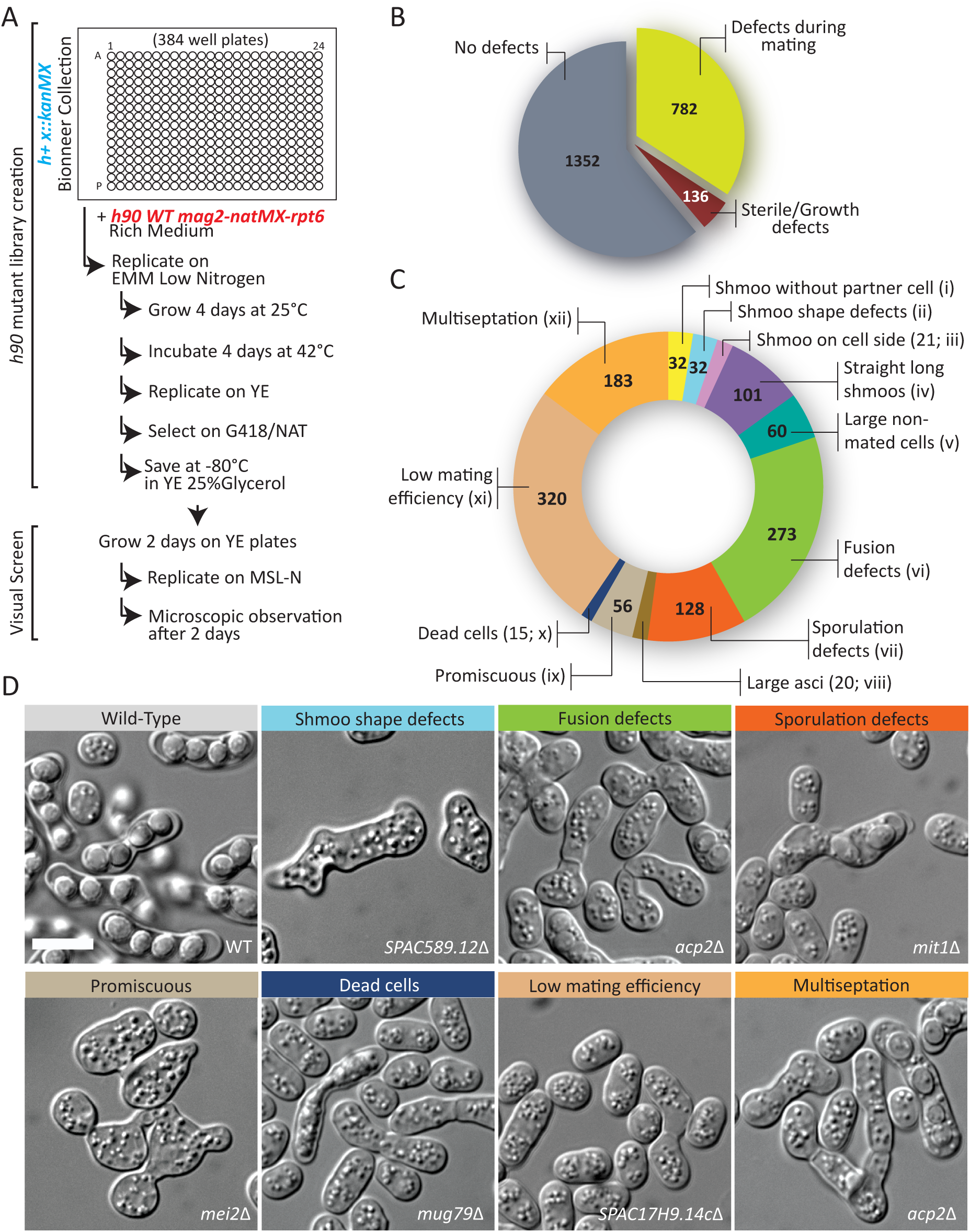
Creation of a homothallic deletion library and visual screening during mating. (A) Workflow used to create the homothallic *h90* deletion library and to visually screen it for morphological abnormalities during mating. (B) Pie chart representing the total number of deletion strains identified to display a visible phenotype during mating. (C)Distribution of all 782 deletion strains with a phenotype in the described phenotypic classes. Note that the total number is >782 because some deletion strains display several phenotypes. (D) Representative images of phenotypic classes. Bars, 5 µm.

From 2270 deletion strains of the *h+* deletion collection, we recovered 2134 *h90* derivatives. The 136 that we could not recover are likely t be either sterile (for example *ste4*∆, *ste6∆*, *ste7∆, ste20∆*, *ras1∆*, *wee1∆*) or too sick to have efficiently crossed, and thus did not give spore progeny in the scheme above. We did not investigate those further at present. To find mutants affecting the mating process, we visually screened the *h90* mutants after a 2-day incubation on solid medium lacking nitrogen(MSL-N). The visual screen was performed in replica by two independent investigators with deletion names undisclosed, in order to eliminate any bias.

Remarkably, out of the2134 screened mutants, 782 mutant showed a visible phenotype during the mating process igure(F 1B). Twelve distinct phenotypes were recorded: these included early mating polarization defects, such as(i) the presence of cells extending growth projections not meeting a partner cell, (ii) aberrant shmoo shapes, (iii) placement, or (iv) length, or (v) the presence of abnormally large unmated cells; (vi) fusion defects, in which paired cells were observed with cell wall at the contact site; and post-fusion phenotypes, such as sporulation defects, in which ascihad abnormal spore numbers o shapes, (viii) abnormally large asci, or(ix) promiscuous cells, in which mutants appeared to mate (or attempt to mate) with multiple partners. We also recorded(x) the presence of dead cells in the mating assay, which may be caused by cell lysis upon deregulated fusion attempts [10], as well as a(xi) *low mating efficiency*class, for mutants in which cell pairs were rare and/or individual cells did not appea be arrested as small cells. Finally, though potentially not starvation specific, we also scored for (xii) multiseptation, in which cells showed multiple septa. For each of these categories, the severity of the phenotype was gauged on a scale from 1 to 10We. note that some deletions were labeled with several distinct phenotypesA. summary of these categories, with the number of identified mutants, represented in Figure 1C. Representative images for some phenotypic classes are shown in Figure 1D. The full description of each phenotypic class, as well as the complete table of mutants with their recorded phenotypes, is available as supplementary material (Supplementary Tables 1 and 2).

### Fusion-deficient mutants

We focused our analysis on the *fusion defects* class of 273 mutants affecting the cell-cell fusion process (Supplementary Table 3). We compared these mutants with a list of genes involved in cell-cell fusion compiled from the literature (Figure 2A). As expected, we identified *fus1∆* and *prm1∆* as fusion defective[9,23]. Deletions of *myo51*, *myo52* and *cfr1* have also been shown to lead to fusion defects[7,13,31], but these strains were absent from the screened library, as were of course deletion of the essential tropomyosin Cdc8 and profilin Cdc3, also required for fusion[11,12]. We also did not identify *dni1∆* and *dni2∆*, likely because these genes are required for fusion only at elevated temperatures [32]. This suggests our screen identified all of the identifiable, previously known genes involved in cell fusio Comparison of our list of fusion-defective mutants also identified several homologues to *S. cerevisiae* cell fusion factors (Figure 2B; see discussion). Amongst all deletions with an arbitrary score of 3 or above, we performed a GO Slim analysis ofthe gene products, which revealed that 12.5% are components of the cytoskeleton, an enrichment relative to the 7.4% withinthe whole genome.

**Figure 2:**
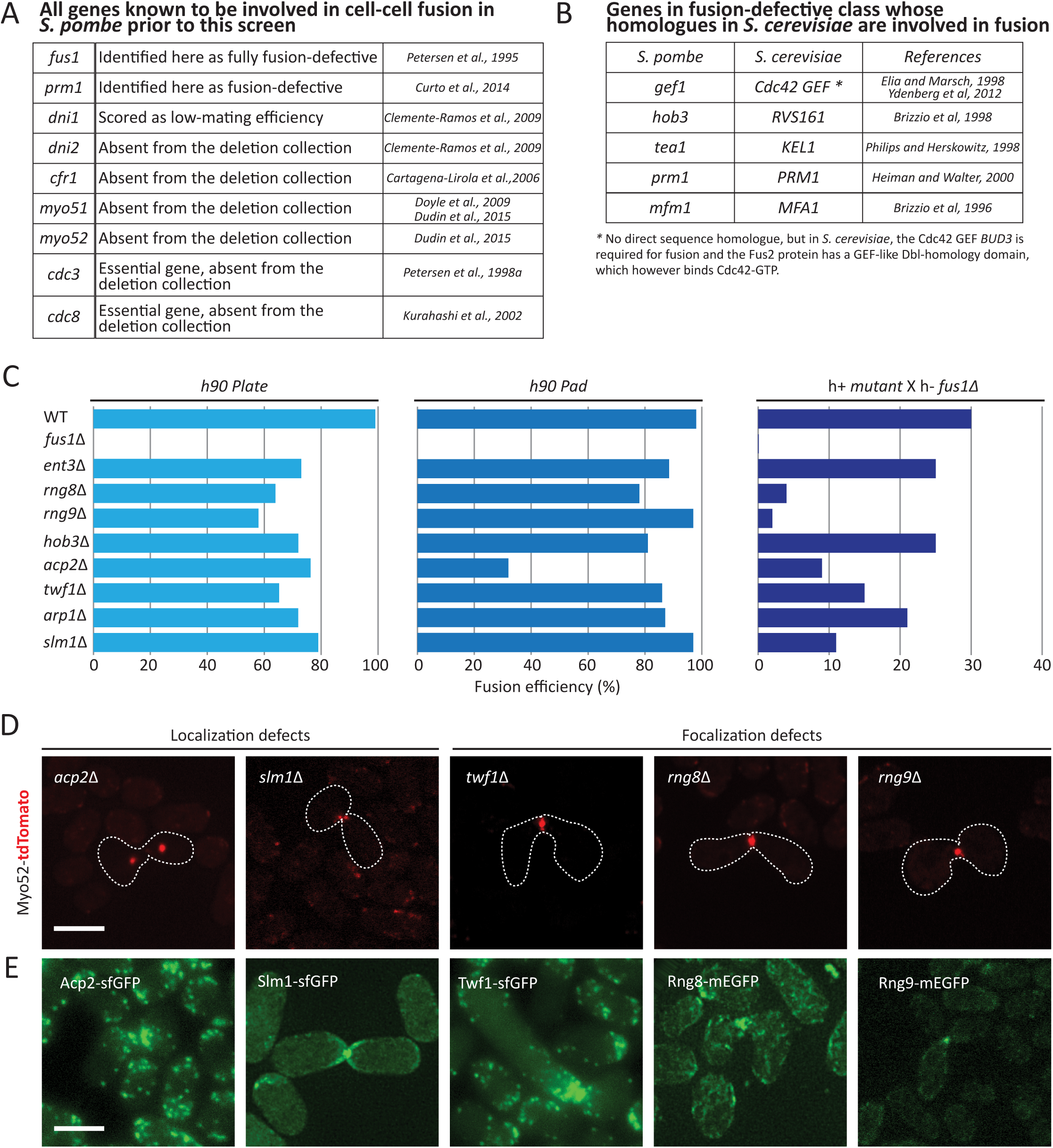
Fusion deficient mutants. (A) List of all *S. pombe*genes known to be involved in cell-cell fusion prior to this screen. (B) List of genes identified to be required for cell fusion with homology to *S.cerevisiae* genes involved in fusion. (C) Fusion efficiency of selected homothallic mutants on plate (left), on a pad between slide and cove-slip (middle) and of heterothallic mutants crossed to *fus1∆* on a pad. (D) Homothallic mutants expressing Myo52-tdTomato. Images shown are maximum intensity projections of a time-series of 7 z-stacks over 15 seconds, except for *slm1Δ*, where a maximum intensity projection of one single time point is shown, which illustrates better the imprecise position of the fusion focus. Compare to wildtype in Figure 3A and 3C. (E) Maximum intensity projection images of *h90* wild-type strains expressing Acp2-GFP, Slm1-mEGF, Twf1-GFP, Rng8-mEGFP and Rng9-mEGFP respectively, during fusionBars, 5 µm.

The enrichment of cytoskeletal components in fusion-defective mutants is interesting becausefusion relies on a dedicatedaster-like actin structure, the fusion focus [7, 10]. To further explore novel fusion-defective mutants affecting the cytoskeleton, we first used publically available data (pombase.org)to discard from the list of cytoskeleton components mutants implicated in chromatin remodeling, spore formation, or with a known localization in the nucleus. This left us with 8 fusion-defective, cytoskeleton-related mutants (Figure 2C), amongst which was the pheromone-dependent formin Fus1 [9]. All 8 strains were verified by PCR for correct deletion of the corresponding gene. The others include the actin capping protein Acp2, previously involved in actin cytoskeleton organization during mitotic growth [33, 34], the centractin family actin like protei Arp1, part of the dynactin complex previously implicated in dynein-dependent nuclear movement during meiotic prophase (horsetail movement; [35]), the actin monomer-binding protein twinfilin *twf1* involved in regulation of polarized growth [36], and the BAR-domain protein Hob3, which was previously known to regulate cytokinesis part through regulation of Cdc42 GTPase [37]. A recently described regulator of the type V myosin Myo51, Rng8, was also selected [21].

To monitor the fusion deficiency of the selected mutants, we used two distinct assays. First, we reproduced the three-dimensional screen conditions by placing cells on SLM-N plates for24h and counting the percentage of non-fused pairs after transfer to a microscope slide (Figure 2C, left). We also used our previously established protocol to quantify fusion efficiency after 24 hours on MSL-N agarose pads, where cells are trapped in a two-dimensional environment for the duration of the sexual reproduction (Figure 2C, middle) [38]. While only *fus1∆* was fully fusion-defective, all mutants showed some fusion defect in at least one of the two assays. We note that, with one exception, the fusion defect was more severe in the three-dimensional assay. This may be due to differences in pheromone distribution or oxygen availability between the two conditions. We also investigated the fusion efficiency on pads of the selected mutants in a heterothallic background with a *fus1* partner, which is fully fusion-deficient. This more stringent test assesses the capacity of the mutant to overcom the fusion deficiency of its partner cellIn. this set-up, again all mutants were more fusion-defective than wildtype cells, with4 mutants highly fusion defective (fusion efficiency < 20%): *rng8*Δ, *acp2*Δ, *twf1* and *slm1* (Figure 2C, right). Deletion of the gene coding for Rng9, the binding partner of Rng8, though not identified in the screen, yielded a similar phenotype as *rng8∆*.

All five deletion strains displayed defects in fusion focus organizatio as labeled with Myo52-tdTomato (Figure 2D): *acp2∆* and *slm1∆* displayed aberrant localization of the fusion focus, with the focus often detached from the cell projection tip in *acp2∆* and out of alignment in *slm1∆*. *rng8∆, rng9∆ and twf1∆* showed wider Myo52- tdTomato signals, suggestive of a defect infusion focus focalization. Fluorescent tagging of each of the five genes at endogenous lo revealed that allaccumulated at the fusion site, albeit with different localization patterns. Acp2 and Twf1 appeared to primarily decorate actin patches as previously shown for Acp2 during vegetative growth Slm1 decorated the cortex of the entire projection tip. By contrast, Rng8 and Rng9 accumulated in a concentrated location at the fusion site (Figure 2D), which coincided with the Myo52-labelled fusion focus (supplementary Figure 1).

Several new genes affecting the fusion process in fision yeast. As all five deletion strains above readily reveal a defect in fusion focus organization, and all encoded proteins localize at the fusion site, we conclude that the screen was highly successful in detecting genes directly involved in the regulation of cell fusion. By extension, this also suggests that many other fusion defective deletion strains will also reveal interesting new cell fusion phenotypes.

### Rng8 and Rng9 are crucial for focalization of the fusion focus

Because Rng8 and Rng9localize to the fusion focus and appear to be required for its focalization, we extended our analysis of their function for the dynamics of the fusion focus during the fusion process, using high-temporal resolution time-lapse microscopy. In *rng8∆*, *rng9∆* and double *rng8∆ rng9∆* mutants mated with wildtype cells, the major fusion focus components Myo52-tdTomato and formin Fus1-sfGFP occupied a zone about twice as wide as in wildtype cells, when measured on sum Z-projections, though the total signal detected at th cell-cell contact site was unchanged (Figure 3A-B). Time-lapse imaging of Myo52-tdTomato in single focal planes further showed that *rng8∆*, *rng9∆* and double *rng8∆ rng9∆* mutants display multiple stable Myo52 dots at the shmoo tip (Figure3C-D). Whereas two dots areoccasionally observed in wildtypecells as the fusion focus forms when the signal matures from a broad crescent-like localization to a single dot (supplementary Figure 2),we never observed several stable dots in wildtype cells. By contrast, in *rng8/9* mutants, most cells exhibited 2, 3 or more dots that were spatially stable over > 1 minute at the cell cortex (Figure 3C-D). This phenotype, as well as fusion efficienc (supplementary Figure 3), were indistinguishable in single and double mutants, consistent with Rng8 and Rng9 forming an obligate dimer [21,22]. Our further analysis was thus conducted only on the *rgn8∆* single mutant. We conclude that the Rng8/9 dimer is required for th formation of a single fusion focus structure.

**Figure 3:**
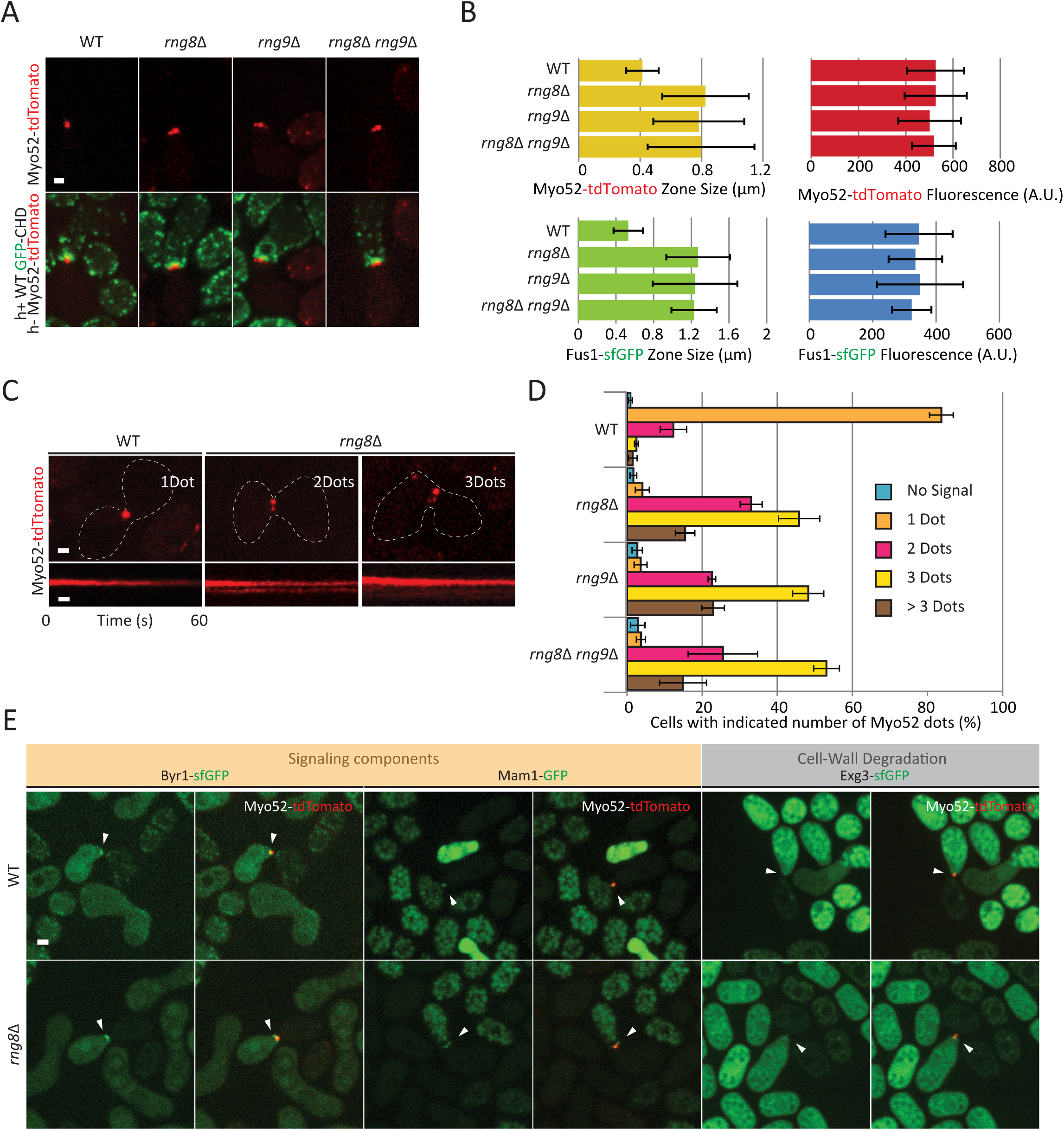
Rng8 and Rng9 are required for the formation of a single fusion actin focus. (A) Cross of wildtype *h+* strain expressing the F-actin marker GFP-CHD with *h-* wildtype, *rng8* Δ*rng9* or double *rng8rng9* mutant strains expressing Myo52-tdTomato, showing a more dispersed Myo52-tdTomato signal in the mutant strains. Images are maximum intensity projection ofa time-series of 7 z-stacks over 15 seconds. (B) Measurements of Myo52-tdTomato and Fus1-sfGFP zone width and fluorescence intensity at the cell-cell contact site. Measurements were done on sum projections of 7-stacksz. (N=15). (C) Typical kymographs of Myo52-tdTomato showing multiple stable dots over >60s in *rng8* in comparison to the unique fusion focus in wildtype. (D) Quantifications of the number of Myo52-tdTomato dots observed in wildtype, *rng8* Δ, *rng9* and *rng8 rng9* mutants. The small percentage of cells showing two or more dots in wildtype likely represents cells captured in the process of forming the fusion foc (see also Supplementary Figure 2)(N>100 cells). (E) Cross of wildtype *h+* cells expressing Myo52-tdTomato with *h-* wildtype or *rng8* expressing Byr1-GFP, Mam1-GFP or Exg3-GFP and Myo52-tdTomato. Images shown are maximum intensity projection ofa time-series of 7 z-stacks over 15 seconds. Arrowheads point at cells in fusion Bars, 2 µm.

Stabilization of the fusion focus relies on accumulation of the pheromone signaling machinery on the structure [10]. In wildtype cells, both M-factor transporter Mam1 and components of the pheromone transduction pathway, including the MAP2K Byr1, strongly accumulate on the fusion focus. In *rng8∆* cells, these components were present at the fusion site, though over a wider region, similar to description of Fus1 and Myo52 above(Figure 3E). Because of weaker signal intensity, we were unable to confidently determine whethere Byr1 and Mam1 also systematically form several distinct stable do or have a more continuous, broad localization, though in some instances, several dots of Mam1 could be clearly distinguished(Figure 3E). This suggests each dot becomes stabilized through the normal pheromone signaling-dependent pathway [10]. This is consistent with the idea that Rng8 is required not for the immobilization of the fusion focus, but for the coalescence ofthe actin aster to a single structure prior to stabilization.

The fusion focus serves for the local release of cell wall diges enzymes [7]. In wildtype cells, the glucanase Exg3-sfGFP can be clearly observed at the fusion focus. In *rng8∆* cells, Exg3 could also be detected at the fusion site(Figure 3E), but only in about half of the cells and often over a wider zone, consistent with the idea that this glucanase is secreted over a broader region upon fusion focus coalescence defects. This defect is consistent with the lower efficiency of *rng8∆* cells in digesting their cell wall, especially when mated wi *fus1∆* partners (Figure 2D). We conclude that the Rng8/9 dimer is critical for the coalescence of the acto-myosin fusion focus into a single aster-like structure, required for local release of cell wall digestive enzymes.

### Rng8 and Rng9 have roles beyond that of regulating Myo51 motor

Previous work has implicated the Rng8/9 dimer in the regulation of the single-headed myosin Myo51. Indeed, Rng8/9 associates with Myo51 in vivo and in vitro and promotes Myo51 cluster formatio Myo51 is not detected on actin cables and only very weakly at cytokinetic ring in *rng8∆* and *rng9∆* cells, and these and *myo51∆* mutants have similar defects in contractile ring assembly [21,22]. One proposed model is that Rng8/9 forms an integral part of the Myo5 motor for most or all of its cellular functions and is strictly required for its processivity [21]. We thus examined in detail the phenotype and localization of Myo51 during cell fusion.

Similar to the situation during cytokinesis [21], Myo51 localization at the fusion focus was strongly reduced, though not completely abolished, in *rng8∆* and *rng9∆* cells (Figure 4A-B). In addition, *myo51∆* cells are partly fusion defective and strongly fusion incompetent when mated with *fus1∆* partners [7], similar to *rng8∆* and *rng9∆* cells. However, in contrast to *rng8∆* and *rng9∆* cells, the Myo52-labelled fusion focus was not significantly broader in *myo51∆* than in wildtype cells (Figure 4C-E). In addition, the vast majority of *myo51∆* cells formed a single Myo52 dot, with only about 30% forming ≥2 dots (Figure 4F). While this is significantly different from the wildtype situation, where about 15% of cells are observed with ≥2 dots, this does not recapitulate the *rng8/9∆* phenotype where about 95% of cells form ≥2 dots. These data indicate that the fusion focus clustering defect of *rng8/9* mutants is not solely due to a loss of Myo51 function Consistently, *rng8∆* and *myo51∆* showed additive phenotypes in fusion efficiency, with the double mutant significantly less fusion-competent than either single mutant. Expectedly, *rng8∆* was also additive with *myo52∆* (Figure 4G). Similar results were consistently observed with the *rng9∆ myo51∆* double mutant (Supplementary Figure 4). Rng8 localization was also significantly broader, though not weaker, at the fusion site in *myo51∆* (Figure 4H-J), suggesting that one role of Myo5 myosin is to concentrate the Rng8/9 dimer in the fusion focus. In conclusion, the myosin V Myo51 and the Rng8/9 dimer each have independent function during fusion and mutually contribute to concentrate the other on the fusion focus.

**Figure 4:**
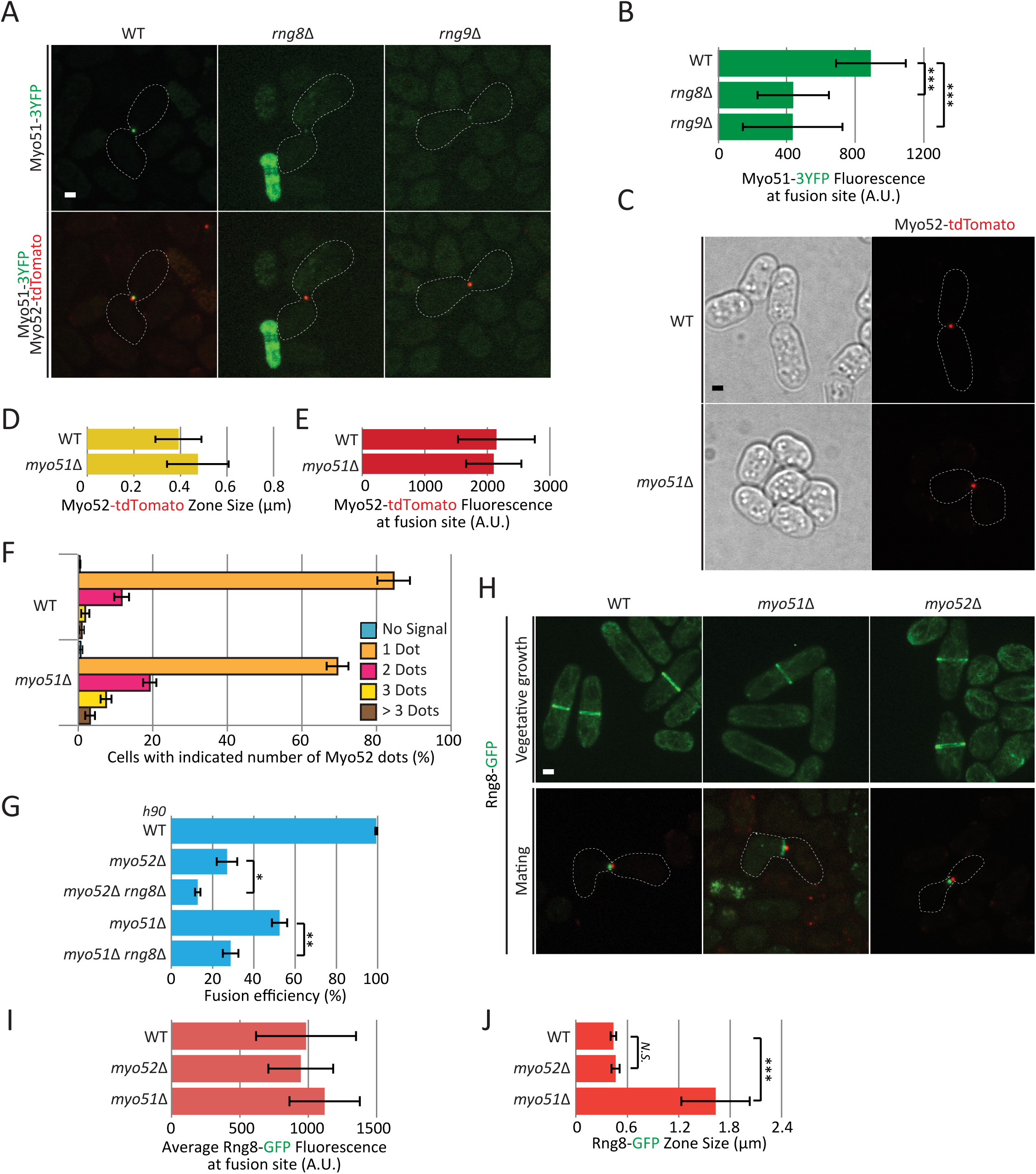
Rng8 and Rng9 have roles beyond that of regulating Myo51 motor. (A) Cross of wildtype *h-* expressing Myo52-tdTomatowith *h+* wildtype, *rng8* or *rng9* strains expressing Myo51-3YFP. (B) Quantification of Myo51-3YFP intensity at the fusion site in wildtype, *rng8* and *rng9* strains as in (A), showing reduction of Myo51 levein the mutants (N>11); (***) P < 5 × 10^−4^, t-test. (C) Localization of Myo52-tdTomato in homothallic wildtype and *myo51* strains. (D-E) Measurements of Myo52-tdTomato zone width (D) and fluorescence intensity (E) at the cell-cell contact sitein strains as in (C)(N=15). (F) Quantifications of the number of Myo52 dots observed in *myo51* mutants in comparison to wildtype (N>100 cells). (G)Fusion efficiency of *h90* wildtype, *myo52* Δ, *myo52 rng8* Δ, *myo51* and *myo51 rng8* strains on pad (N>200); (*) P < 0.04, (**) P < 0.003 t-test. (H) Localization of Rng8-mEGFP in *myo51∆* and *myo52∆* mutants during vegetative growth (top) and during mating (bottom) in *h+* cells crossed to a wildtype *h-* strain expressing Myo52-tdTomato. (I) Measurements of average Rng8-mEGFP fluorescence at fusion site in strains as in H (N=10)(J). Measurements of Rng8-mEGFP total zone width at the cell-cell contact sitein strains as in (H) (N=10); (***) P < 7 × 10^−5^, t-test. Bars, 2 µm.

### A tropomyosin point mutant recapitulates the *rng8/9∆* phenotype

Recent in vitro work has shown that the Rng8/9-Myo51 complex binds tropomyosin-decorated actin filaments independently of the Myo51 motor domain [22]. This binding was proposed to anchor the complex to tropomyosin-decorated filaments to favor their transport along other actin filaments bound by the motor domainThis. prompted us to examine the role of tropomyosin Cdc8in actin focus formation.

Cdc8 was previously shown to be necessary for cell fusion and to localize at the fusion site [11]. Cdc8-GFP, expressed under the inducible *nmt41* promoter [39], indeed accumulated at the fusion sit in both wildtype and *rng8∆* cells to similar levels, though it occupied zone about twice as wide in *rng8∆* cells, as described above for other fusion focus components(Figure 5A-C). We also confirmed that cells of the temperature-sensitive *cdc8-382* mutant [40], though able to form pairs, were highly fusion-deficient at the semi-permissive temperature of 33ºC (Figure 5D). These data confirm an important role of tropomyosin in cell fusion.

**Figure 5:**
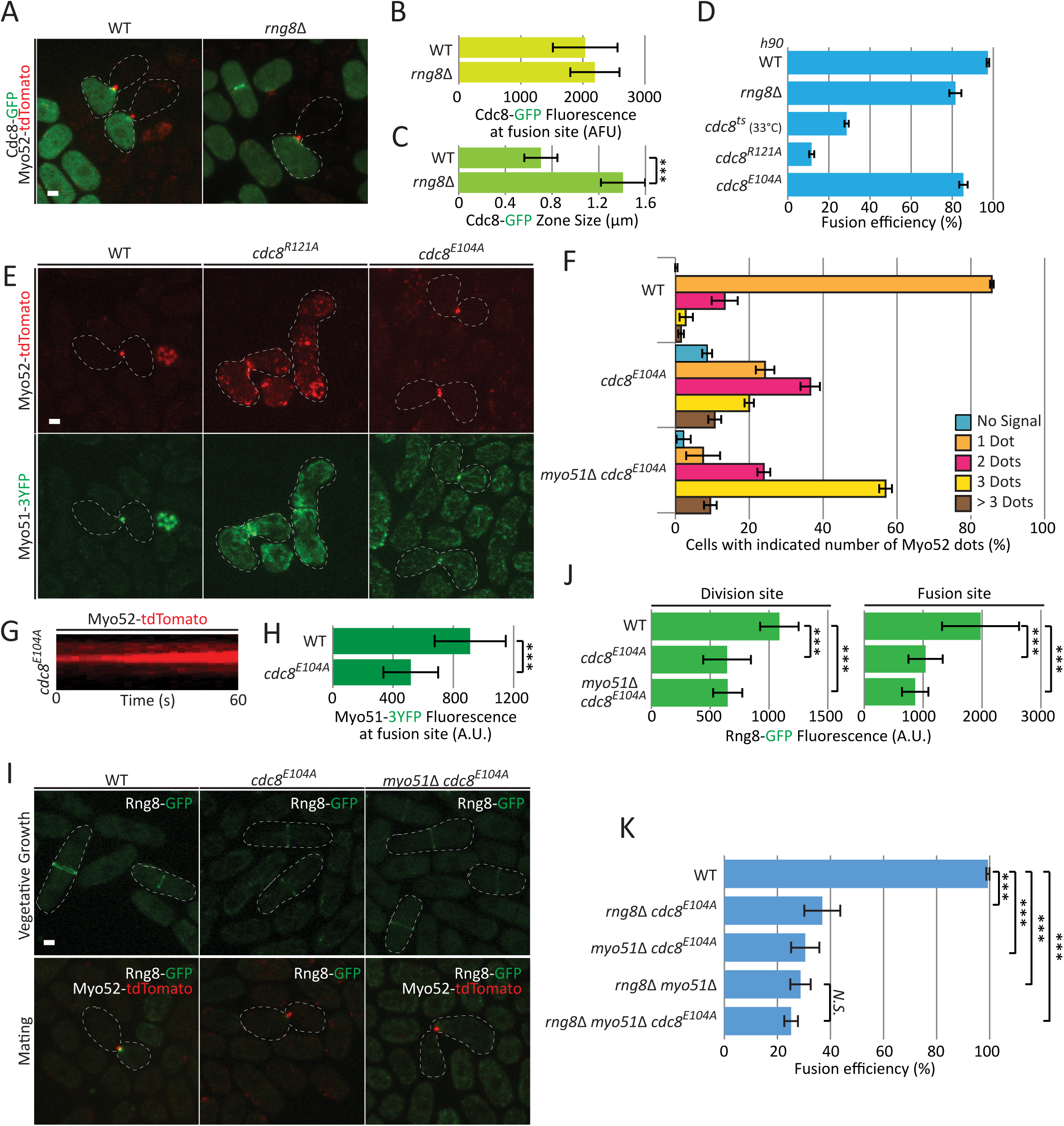
A tropomyosin point mutant recapitulates the *rng8/9* deletion phenotype. (A) Cross of wildtype *h+ myo52-tdTomato* with *h*-wildtype or *rng8* cells expressing nmt41-Cdc8-GFP. (B)Measurements of Cdc8-GFP accumulation at fusion site (N=15).(C) Measurements of Cdc8-GFP zone width at the cell-cell contact site (N=10); (***) P < 10^−7^, t-test. (D) Fusion efficiency of *h90* wildtype, *rng8* Δ, *cdc8-382, cdc8*^*R121A*^ and *cdc8*^*E104A*^ mutants on pads (N>600). Note that allstrains were incubated at 30ºC except for *cdc8-382*, which was incubated at the semi-restrictive temperature of 33ºC. (E) Localization of Myo52-tdTomato and Myo51-3YFP in *h90 cdc8*^*R121A*^ and *cdc8*^*E104A*^ mutants. (F) Quantifications of the number of Myo52-tdTomato dots observed in *cdc8*^*E104A*^ and *cdc8*^*E104A*^ *myo51Δ* mutants in comparison to wildtype (N>100 cells). (G)Kymographs of Myo52-tdTomato showing multiple stable dotsover >60s in *cdc8*^*E104A*^ mutant. (H) Myo51-3YFP fluorescence at the fusion site is decreased i *cdc8*^*E104A*^ mutant (N=15); (***) P <3 × 10^−5^, t-test. (I) Localization of Rng8-mEGFP in *h+ cdc8*^*E104A*^ and *myo51cdc8*^*E104A*^ during vegetative growth and when crossed to *h-* Myo52-tdTomato wildtype cells during mating. (J) Measurements of Rng8-mEGFP fluorescence at the division and the fusion sites show a significant reduction in the *cdc8*^*E104A*^ and *myo51cdc8*^*E104A*^mutant allele (N= 15);(***) P <1.7 × 10^−8^, t-test. (K) Fusion efficiency of wildtype, *rng8cdc8E104A, myo51cdc8E104A, rng8 myo51cdc8*^*E104A*^ and *rng8 myo51* (N>150); (***) P <6 × 10^−5^, t-test. Bars, 2 µm.

We then took opportunity of a collection of point mutations predicted surface-exposed Cdc8 residues conserved in fungi [41, 42] to screen for non-conditional mutants that would hinder cell fusion when homothallic. This identified two alleles each carrying a single point mutation,*cdc8*^*R121A*^ and *cdc8*^*E104A*^, with reduced fusion efficiency (Figure 5D). The phenotype of *cdc8*^*R121A*^ was very severe, with only about 10% fusion efficiency. The *cdc8*^*R121A*^ mutation causes significant actin cytoskeleton organization defects during vegetative growth including weak actin cables, dispersed patches and defective cytokinetic ring, and reduces the affinity of tropomyosin for actin about 30-fold in vitro [42]. During mating, both Myo52-tdTomato and Myo51-3YFP failed to concentrate at a single focal point at the fusi site in this strain, though they were enriched at the zone of cell-contact, strongly suggesting that the global organization ofthe actin cytoskeleton is affected and the fusion focus does not form. We conclude that the fusion defect observed in *cdc8*^*R121A*^ cells is due to strongly reduced actin-tropomyosin interaction.

The second fusion-defective allele, *cdc8*^*E104A*^, displayed a much milder fusion problem, similar to that observed in *rng8∆* (Figurer 5D). This allele was shown to have some very mild cell polarization defects during vegetative growth, but does not affect actin binding [42]. Remarkably, the localizations of Myo52 and Myo51 during mating strongly resembled those observed in *rng8∆* cells: most *cdc8*^*E104A*^ cells exhibited 2 or3 Myo52 dots that were spatially stable at thecell-cell contact site (figure 5E-G), though we note the phenotype was not quite as severe as that of *rng8∆* (see Figure 3D). In addition, Myo51 was present in significantly reduced amounts (Figure 5H). The similarity of the *cdc8*^*E104A*^ and *rng8∆* phenotypes suggest that *cdc8*^*E104A*^ affects Rng8/9 function. Indeed, Rng8 was strongly delocalized from all actin structures: it could not be detected on actin cablesand only weaklyon the cytokinetic ring during vegetative growth, as well as onthe fusion focus during mating (Figure 5I-J). As observed in wildtype background, we note that this localization was not further weakened by deletion of Myo51 (Figure 5I-J). These results suggest that the fusion defects observed in *cdc8*^*E104A*^ stems from a loss of binding with the Rng8/9 dimer.

Two pieces of data suggest that interaction of the Rng8/9 dimer with both tropomyosin and myosin V Myo51 contribute to focalization of the fusion focus. First, construction of a double mutant *cdc8*^*E104A*^ *myo51∆* exhibited more severe de-clustered focus phenotype than either single mutant, identical to *rng8∆* (compare Figure 5F and 3D). Second, epistasis analysis showed that the triple *cdc8*^*E104A*^ *myo51∆rng8∆* mutant was not more fusion-defective than the double *myo51∆ rng8∆* mutant, suggesting the *cdc8*^*E104A*^ mutation does not affect other components than Rng8 and Myo51 (Figure 5K). By contrast both *cdc8*^*E104A*^ *rng8∆* and *cdc8*^*E104A*^ *myo51∆* double mutants were significantly more fusion-defective than the corresponding single mutants (Figure 5K, compare to Figures 4G and 5D), suggesting the *cdc8* mutant weakens the interaction of both Rng8/9 and Myo51 with actin filaments sufficiently to abolish the function of the complex. We conclude that Rng8/9 acts through both tropomyosin and myosin V Myo51 to organize the fusion focus.

## Discussion

### The homothallic deletion collection: a new genetic tool

Systematic gene deletion collections in both budding and fission yeasts have enabled important advances in the understanding fundamental cellular processes [28,43,44]. To facilitate the discover of genes with function in the sexual reproduction process, we derived a self-fertile (homothallic) version of the collection of viable deletions in fission yeast. Because both partner cellscarry the same deletion this approach is more sensitive in identifying genes important for the mating process, whose presence in one of the two partners may be sufficient for functionality. This allowed the discovery of>200 genes involved in cell-cell fusion, a process previously noted for its robustness [4].

This approach also ensured that diploid zygotes were homozygote mutant, leading to the discovery of sporulation-deficient mutants. A similar strategy, using a homothallic derivative of the deletion collection to screen for sporulation-defective mutants through absence of iodine staining, which specifically stains spores, was published during the course of our work [27]. Our list of sporulation-defective mutants overlaps with that described in this work, but is more extensive (Supplementary Tables 4 and 5), likely because visual screening permitted identification of more subtle phenotypes, for instance of abnormal spore number.

Besides these two large phenotypic classes, a large number of deletions strains were identified with defect in cell polarization, and categorized in several phenotypic classes. Cell polarization in response to pheromone, leading to cell-cell pairing, is a complex process involving an exploratory patch of active Cdc42 GTPase that serves as site of pheromone release and signaling [5,6]. We note that genes involved in cell polarization during vegetative growth, though present in the deletion collection, were not prominent the *shmoo shape defects* class, suggesting that regulatory mechanisms of polarized growth are in part distinct, as also previously suggested [45] Mutants with aberrantly placed shmoos, absent from cell sides, or formed in absence of a partner maybe caused by a defect in the Cdc42 exploratory polarization mechanism, or may reflect an alteration in pheromone signaling or perception, whichmodulates exploratory polarization [5, 6].

Finally, one unexpected and very interesting category of mutants is the *promiscuous* class. While wildtype cells always atem with a unique partner, yielding diploid zygotes, these mutants showed multiple cell projections to several partner. While time-lapse microscopy will be required to ascertain whether cells shmoo in all direction atthe same time or sequentially, and whether they fuse or only attempt to with several partners, we confirmed that deletion of the master regulators of meiosis *mei2* and *mei3* [46–48] show successive fusion with multiple partners. This phenotype was so extensive in the screen that asci were not readily identified and thus the absence of spores was missed The. mere existence of this category of mutants indicates the existence of regulatory mechanisms that arrest mating in zygotes and thus ensure the alternance of haploid and diploid generations (A. Vjestica, LM and SGM, manuscript in preparation).

In summary, our visual screening of a homothallic derivative of the collection of viable gene deletions exposes a host of novel gene functions that begin revealing new biology and provides a rich basis for future research. This homothallic deletion collection also represents a novel resource that can be further screened for more specific phenotypes.

### Fusion-deficient mutants: commonalities for fusion in ascomycetes

We focused on the class of fusion-defective mutants, which represents the largest well-defined phenotypic class. The identified mutants may affect any of the multiple steps required to achieve cell-fusion, from signaling, cell-cell adhesion, cytoskeletal organization, cell wall digestion to plasma membrane fusion. We note that no other deletion than *fus1∆* showed a fully penetrant phenotype. This may be due to three main reasons. First, there is significant redundancy between components and/or pathways, as also noted in the study of-cell fusion in budding yeast and *Drosophila* myoblasts [4]. For instance, neither Myo51 nor Myo52 is essential for fusion, yet double deletions fully abrogate it [7]. Second, some components may be re-used several times during the mating process, such that their deletion blocks mating at an earlier stage than fusion. This is for instance the case the pheromone-MAPK cascade, essential for sexual differentiation, but which re-localizes to the fusion focus to signal fusion commitment [10]. Finally, fusion may rely on components otherwise essential for viability, which could not be identified in this screen. For instance, fusion requires a dedicated actin structure, the fusion focus, which besides its formin nucleator Fus1, is built from components also necessary during cell division [7, 11, 12]. However, this screen provides a very large entry-point into the fusion process.

It is interesting that the homologues of several genes or pathways required for cell fusion in *S. cerevisiae* were identified as fusion defective in our screen (Figure 2B). These include in particular the BAR adaptor Hob3, which binds the Cdc42 guanine nucleotide exchange factor Gef1 andhelps promotes GTP exchangeon Cdc42 [37], and Gef1 itself. The *S. cerevisiae* homologue of Hob3, Rvs161p, regulates fusion through interaction with Fus2p [49]. While Fus2p has no identifiable sequence homolog in *S. pombe*, it directly binds active Cdc42p, and both Cdc42p and its guanine exchange factor Cdc24p are required for fusion [50,51]. The Cdc42 GEF Bud3p also contributes to cell fusion in *S. cerevisiae* [52,53]. We also found that the PAK kinase Shk2 is required for fusion, arguing that a common set of proteins around Cdc42 regulates cell fusion in both organisms Similarly,. the deletions of Tea1 and Tea4, important regulators of cell polarity delivered to cell poles by microtubules during mitotic growth [54–56], are present in the fusion-defective class. In *S. cerevisiae*, the homologue of Tea1, Kel1p, promotes cell fusion through regulation of Fus2p localization [57,58]. Finally, we identified one of the M-factor coding genes *mfm1* in the fusion-defective class. This is consistent with the notion that fusion commitment in *S. pombe* requires a sharply graded pheromone signal [10], and similar to findings *S. cerevisiae* where repression of *mfa1*, coding for *a*-factor, or mutation of its transporter lead to cell fusion defects [59,60]. Finally, as noted previously, formin activities (Fus1 in *S. pombe* and likely Bni1 in *S.cerevisiae*) and the multi-pass trans membrane protein Prm1 are required for fusion in both species [4,9,23,24, 61]. Together, these findings suggest that the process of cell-cell fusion is likely to be highly conserved between these two distant ascomycete species.

### Rng8 and Rng9 are required for fusion focus clustering before stabilization

The phenotype of *rng8∆* and *rng9∆* is distinct from previously reported phenotypes: the fusion focus is partly de-clustered, yet each dot is spatially stable and appears to accumulate pheromone-signaling components. Formation of the fusion focus in wildtype cells initiates from a broad distribution of Myo52 at the cell projection cortex, which coalesces into a single focus [7]. Intermediate multi-dots stages resembling the *rng8∆* phenotype can be transiently observed, but the small clusters are not maintained over time and immobilization happens only for a single structure. We suggest that Rng8/9 normally acts before fusion focus stabilization to ensure the formation of singular actin aster.

The outcome of the de-clustered focus is that cell wall digestiv enzymes are not released at a single location. In the wildtype situation, cell wall hydrolytic enzymes (glucanases) are released specifically at the fusion focus, while glucan synthases are broadly localized, yielding aprobable gradient of cell wall hydrolytic activity [7]. When the fusion focus is de-clustered, this gradient likely cannot be well established. Consistently, the glucanase Exg3 was difficult to detect. The consequence is that in crosses to *fus1∆*, *rng8∆* cells are largely unable to overcome the homogeneous release of hydrolytic enzymes by their partner, and thus fusion fails. By contrast, when mated to wildtype or itself, *rng8∆* cells often succeed in cell wall digestion, likely because there is one dominant focus.

### Rng8/9 clusters the fusion focus through tropomyosin interaction

The Rng8/Rng9 complex has emerged as an important regulator of the type V myosin Myo51 [21,22]. Myo51 is an unusual myosin V: in vitro work has shown it is largely monomeric, has a low duty-ratio and is unable to move continuously on actin as a single molecule [22,62]. Similarly, dim punctae of Myo51 thought(to represent dimers)do not move processively on actin cables in vivo [21]. However, assemblies of several Myo51 molecules display processive movements both in viv and in vitro. Two hypotheses have been proposed for the role of Rng8/9. Rng8 and Rng9 co-purify as oligomers from cells and these proteins convert non-processive Myo51 punctae into processive larger assemblies in vivo. Thus a first model is that Rng8/9 converts Myo51 into a processive motor through cluster formation [21]. Recent in vitro work has shown that Rng8/9 also provides an ATP-independent binding site for the Myo51-Rng8/9 complex to bind tropomyosin decorated actin, independently of the Myo51 motor domain. This immobilizes the complex when bound to a single filament, but promotes filament bundling or sliding, depending on assay conditions, when two distinct filaments are connected [22]. Thus, a second hypothesis is that Rng8/9 anchors Myo51 to a neighboring actin filament, in a tropomyosin-dependent manner, to favor filament bundling and/or sliding.

Our data lend strong in vivo support for the importance of the Rng8/9-tropomyosin interaction in the assembly of the fusion focus. Tropomyosin was known to be critical for cell fusion [11], and we have confirmed, through use of the *cdc8*^*R121A*^ allele, which displays 30-fold lower actin binding [42], that tropomyosin-actin binding is indeed essential. We now show that *rng8* deletion and *cdc8*^*E 1 0 4A*^, a tropomyosin point mutant that strongly compromises Rng8 localization to actin structures but does not affect actin binding (our data and [42]), yield almost indistinguishable phenotypes in fusion focus de-clustering. These data predict that the highly conserved region around tropomyosin E104 [42] serves as specific binding site for Rng8/9, though this will need to be confirmed through in vitr re constitution studies. We note that the additive phenotype of the *rng8∆ cdc8*^*E104A*^ double mutant suggests that this region on tropomyosin also plays a role in Myo51 binding. Because Rng8 localization to actin structures was also strongly compromised in *cdc8*^*E104A*^ vegetative cells, it will be interesting to investigate the possible cytokinetic defects and epistasis of *cdc8*^*E104A*^ in comparison to *rng8∆*, to generalize these findings to all actin structures.

By contrast, the interaction between Rng8/9 and Myo51 appears less critical for fusion focus organization. Myo51 likely plays a small role but its deletion shows only very weak declustering phenotype and is strongly additive to *rng8∆* in terms of fusion efficiency In. addition, the observation that Rng8 fails to be enriched on the fusion focus in *myo51∆* cells suggests the prime function of the Myo51-Rng8/9 interaction during fusion may be to concentrate Rng8/9 at the fusi site. We conclude that Rng8/9 binding to tropomyosin-decorated actin is critical to focus the actin fusion structure.

The fusion focus may be in some ways considered an analogous actin based structure to the microtubule-based mitotic spindle pole. Spindle pole focusing strongly depends on minus-end directed motor proteins [63], but also of non-motor microtubule-associated proteins. In particular,the non-motor spindle matrix protein NuMA has activities very analogous to those of Rng8/9 in spindle pole focusing: NuMA forms dimers or oligomers, and binds both the pole-directed dynein complex and microtubules directly. Thus, it may focus spindle poles through two possible scenarios: by forming a dynein-NuMA complex that provides two MT binding sites to cross-link and slide MTs passed each other or through NuMA oligomers that directly cross-link MTs [64–66].

Our data suggests that the Rng8/9 complex functions in the fusio focus much like NuMA at the spindle pole With. membrane-proximal Fus1 nucleating actin filaments that are decorated by tropomyosin, Rng8/9-tropomyosin interaction may promote filament-filament interactions and focus formation in two complementary ways. Formation of a complex with Myo51 may allow concentration of Rng8/9 and sliding of filaments (as proposed in [22]) towards the membrane-proximal barbed end. This would contribute to the coalescence of actin filaments to a single focal point, though our data suggest this contribution is modest. Alternatively, and likely more prominently, Rng8/9 may form oligomeric assemblies that crosslink tropomyosin-decorated actin filaments in absence of motor. As oligomers were not detected in vitro [22], their formation may be indirect or require specific post-ranslational modification. Cross-linking of filaments may selectively stabilize these filaments, thus leading to progressive structure focalization.The Rng8/9-dependent mode of fusion focus clustering may represent one of several mechanisms. Future study of the here-identified collection of deletion promises to reveal fundamental mechanisms of cytoskeletal organization and cell fusion.

## Materials and Methods

### Yeast strains and culture

Strains used in this study are listed in Supplementary Table 6. For assessing exponentially growing cells, cells were grown in Edinburgh minimal medium (EMM) or minimal sporulation media with nitrog (MSL+N) supplemented with amino acids as required. For assessing mating cells, liquid or agar minimal sporulation media with nitrogen (MSL-N) were used [38, 67]. All live-cell imaging was performed on MSL-N agarose pads [38]. Mating assays were performed as in [5, 7, 38]. Briefly, pre-cultures of cells were grown at 25°C OD600 = 0.4-1 in MSL + N (for heterothallic crosses, cellswere mixed in equal parts), diluted and grown for −1820 h to OD600 = 0.4– 0.6 at 30°C in MSL + N. Cells were pelleted by centrifugation and washed three times in MSL-N and mounted onto MSL-N 2% agarose pads and sealed with VALAP. Pads were then incubated for either 1 h at 25° before imaging in overnight movies or overnight at 18°C before imaging. Fusion efficiency was measured as in [7, 10].

### Genetic Screen

The haploid *S. pombe* deletion mutant library was purchased from Bioneer (South Korea). The deletion strains are marked with a G418-resistance *kanMX* cassette in an *h+* strain background (*h+ ade6-M210ura4-D18 leu1-32*). To examine phenotypic changes during mating, an *h90* library was created by crossing the collection of deleted mutants with a homothallic strain carrying a Nat-resistance *natMX* cassette at the *h90* locus (YSM2945 *h90 mag2-natMX-rpt6*). This strain was made by a PCR based approach using primers osm1023-(5’-<u>caacaagagctgcgttgactgctttttttttgctatataatccagatgcagattattttaaaatactaatccaaatat</u>CGGATCCCCGGGTTAATTAA) and osm1024 (5’<u>ttaatgggttgtttgtcagtcgttgatttagtcctgaatatacataaggaaaagttaatccagggtggagtcgactct</u>GAATTCGAGCTCGTTTAAAC-) to amplify the natMX cassettfrom pFA6a-NatMX6. This product was designed to recombine into the intergenic region between the mag2 and rpt6 open reading frames the mat locus(homology is underlined).

Before performing the phenotypic analysis, the collection was amplified and frozen down at-80°C. For amplification, the deletion strains were inoculated in200µl MSL+N in 96-well plates with the help of a Tecan® robot and incubated at 30°C with shaking for 2 days. Pre-cultures of *h90 mag2-natMX-rpt6* cells were grown at 25°C to OD600 = 0.4–1 in MSL+N, diluted and grown for 18–20 h to OD600 = 0.4 at 30°C inMSL+N. 25µl of each deletion strain cultures were mixed with 25µl of the *h90* strain in a 96-well plate and 2µl of the mixture were spotted on EMM-ALU plates containing low nitrogen amounts (24mM NH4Cl) with the help of a Tecan® robot. Plates were incubat at 25°C for 4 days to allow mating and sporulation, and then shifted to 42°C for 3 days to kill un-sporulated diploid and un-mated haploid cells. The EMM-ALU plates were replica plated on YE with the help o Singer® robot and incubated for 2 days at 30°C to allow spore germination and colony growth. YE plates were replica plated on YE plates containing both G418 and nourseothricin (250µg/ml G41 100µg/ml Nat) and incubated for 2 days at30°C to select for *h90* deletion strains. To freeze down the collection, mutants were inoculated from YE-G418/Nat plates in 200µl YE in 96-well plates with the help of a Tecan® and a Singer® robot. Cells were grown30°Cat for 2 days and100µl YE containing 50% glycerol was added with th help of a Tecan® robot before freezing down the strains at −80°Cat.

For phenotypic analysis of *h90* deletion strains, the homothallic mutants were first spotted on YE and growth at 25ºC for 2 days, then replica plated on MSL-N and incubated at 25ºC for 2 day Mutants were visually screened on a small table-top Leica microscope with 40x magnification for mating defects. Practically, cells were picked up with a toothpick and resuspended in 2µl MSL-Non a glass slide and coated with a coverslip. The analysis was done in duplicatby 2 independent investigators and phenotypic defects were classified and scored between 1 and 10. Mutants with score ≥ 5 were screened second time to confirm the phenotypic defectWe. note that diploid killing during *h90* collection generation was largely efficient as we observed a few azygotic tetrads (issued from the sporulation of a diploid, rather than a freshly formed zygote) in only 76 strains through the entire visual screen. GO enrichments were performed using GO term finder (http://go.princeton.edu/cgi-bin/GOTermFinder

### Microscopy and Image Analysis

The spinning-disk microscope system, previously described [16] was used throughout the study. Optical slices were acquired every 0.6 µm and all panels show maximum projections, unless otherwise indicated For zone size measurements, fusion efficiency and number of Myo5 dots at fusion site (Figures 2D, 3B, 3D, 4B, 4C, 4F, 4I, 5C, 5D, 5F and 5K), the plugin ObjectJ in ImageJ (National Institutes of Health) was used. Fluorescence intensities of Myo51-GFP, Rng8-GFPand nmt41- cdc8-GFP in Figures 3B, 4D, 4G, 5B, 5I and 5J were measured in ImageJ using a manually drawn area around the shmoo tip in maximum projections of seven slices over −4µm total depth. Background fluorescence was measured and subtracted from eth original measurements. Kymographs in Figures 3C and 5H were constructed in Image J version 1.47 (National Institutes of Health) by a drawing 3 pixel-wide line at the cell tip Figures. were assembled with Adobe Photoshop CS5 and Adobe Illustrator CS5. All error bars are standard deviations. All experiments were done a minimum of three independent times, and statistical analysis was done across repeats of the same experiment.

## Author contributions

LM and FB derived the *h90* deletion collection and performed het visual screen, with the help of VV. OD conducted the secondary screen with the help of RG, and performed all other experiments. SGM supervised the study and wrote the manuscript with contributions from OD, LM and FB.

## Acknowledgements

We thank Bart Deplancke (EPFL) for use of the Tecan and Singer robots, Sarah Hitchcock-DeGregori (Rutgers University) for the *cdc8* hypomorphic alleles and James Moseley, Jian-Qiu Wu and David Kovar for strains. This work was supported by an ERC Consolidator grant (CellFusion) and a Swiss National Science Foundation (31003A_155944) to SGM.

